# Photonic-Plasmonic Coupling Enhanced Fluorescence Enabling Digital-Resolution Ultrasensitive Protein Detection

**DOI:** 10.1101/2022.10.29.514360

**Authors:** Priyash Barya, Yanyu Xiong, Skye Shepherd, Rohit Gupta, Lucas D. Akin, Joseph Tibbs, Han Keun Lee, Srikanth Singamaneni, Brian T. Cunningham

## Abstract

Assays utilizing molecular fluorophores are common throughout life science research and diagnostic testing, although detection limits are generally limited by weak emission intensity, thus requiring many labeled target molecules to combine their output to achieve signal-to-noise greater than the background. Here, we describe how the synergistic coupling of plasmonic and photonic resonance modes can significantly boost the emission from fluorescent dye molecules without increasing the illumination intensity while utilizing a microscopy approach with a broad field of view. By optimally matching the resonant modes of a plasmonic fluor (PF) nanoparticle and a photonic crystal (PC) surface with the absorption and emission spectrum of the PF’s fluorescent dye, we observe a 52-fold improvement in signal intensity, enabling individual PFs to be observed and digitally counted, using an approach in which one PF tag represents detection of one target molecule. The photonic amplification from the PF can be attributed to the strong near-field enhancement due to the cavity-induced activation of the PF, PC band structure-mediated improvement in collection efficiency of emitted photons, and increased rate of spontaneous emission. We demonstrate the applicability of the method by dose-response characterization of a sandwich immunoassay for human interleukin-6, a biomarker commonly used to assist diagnosis of cancer, inflammation, sepsis, and autoimmune disease. We achieve a limit of detection of 10 fg/ml, representing a capability three orders of magnitude lower than standard immunoassays.

## INTRODUCTION

Fluorescence-based bioanalytical techniques are among the most predominant methods for quantitative detection of many classes of biomolecules^1-3^. The basic working principle involves attachment of the fluor to a biomolecule with a covalent chemical bond, and with knowledge of the absorption and emission spectra of the fluor, selecting an appropriate illumination source and emission filter to enable observation with a single-element photodiode or an array of photodiodes in an image sensor. Fluorescent dyes are commonly utilized elements within flow cytometers, microscopy, lab-on-a-chip systems, and health diagnostics instruments due to their versatility in terms of selective attachment to other molecules, low cost, wavelength versatility, and ease of use. However, the main limitations for fluorescent dyes in the context of bioassays are rapid photobleaching, and the low number of detectable photons/fluor, as dictated by fluorescent lifetime, collection efficiency, and excitation efficiency. Typically, an efficient photon-collecting high numerical aperture (NA) objective or electron-multiplying charge-coupled device camera is required for imaging single fluorescent tags^2, 4^, resulting in expensive instrumentation, that limits their use in point-of-care diagnostic testing and routine life science research. When individual fluorophores are used to label individual target biomolecules in an assay, the limitation manifests itself in the requirement to aggregate many analytes together, so as to combine the output of many fluors. Many emitters are required to enable observation above noise due to the combined effects of background fluorescence from the environment and the inherent noise of the photodetector (dark current and shot noise). Thus, detection of target biomolecules in fluorescent assays on ordinary surfaces, such as glass and plastic, typically achieve detection limits in the 1-100 pg/ml range as observed in array formats in which a capture molecule is applied to an assay surface in the form of a small spot ^5-7^. Prior research in the use of resonant dielectric nanostructures, such as photonic crystals (PCs)^8-10^, zero-mode waveguides^11, 12^, microring resonators^13^ and plasmonic surfaces^14, 15^ have sought to overcome the inherent brightness limitations of fluorophores through enhancement in the excitation intensity, increased collection efficiency of emitted photons through directional extraction, and reduced lifetime through the Purcell effect.

While these approaches have proven effective for reducing detection limits for biosensing assays, the techniques are not capable of reducing limits of detection and quantitation to the ultimate limit of digital resolution, in which each detected molecule may be counted individually^16, 17^. While digital resolution biodetection may be achieved through partitioning of the sample into thousands of individual droplets, followed by enzymatic amplification, as with ddPCR^18^, BEAMing^19^, and Quanterix Simoa™ assays^20^, we seek a more direct and simple approach in which a nanoparticle tag is attached to each target molecule, and a simple and inexpensive optical detection instrument counts the tags with high signal-to-noise ratio.

Plasmonic nanoparticles and nanoantennas produce near-field fluorescent enhancements due to their small modal volumes^21-23^ while also drastically increasing the local density of states (LDOS) near the emitter, thus increasing the rate of spontaneous emission^24, 25^. Recently, by coating a plasmonic nanostructure with fluorophores, stable and ultrabright fluorescent nanoconstructs have been applied as fluorescent reporter tags^26, 27^. However, plasmonic nanostructures are also associated with high losses due to nonradiative processes and lack of directive emission^28, 29^. One approach to mitigate the losses has been coupling the plasmonic nanoantenna to a photonic resonator which could effectively depolarize the antenna, lowering its absorption in a narrow spectral window^30, 31^. Additionally, the resonant matching of these two oscillators can result in an enormous increase in near-field enhancement^26, 32^. Utilizing a dielectric structure like a photonic crystal (PC) could also mitigate the lack of directive emission by enabling controlled Bragg scattering^8, 33^.

In this work, we describe an approach for achieving digital-resolution detection of target biomolecules through the synergetic interplay of plasmonic and photonic resonances to boost the emission of fluorescent dye molecules. The approach builds upon our recent exploration of plasmonic-photonic hybrid coupling between plasmonic nanoparticles and a PC surface, in which enhanced absorption cross section results from strategic selection of the nanoparticle LSPR wavelength with respect to the PC band structure^30^, which we demonstrated as an effective means for enhanced surface-enhanced Raman scattering^32^ and electromagnetically enhanced catalysis of chemical reactions^16^. The key benefits the hybrid coupling brings is (1) the strong near-field enhancement which tightly focuses the excitation energy for the absorption by fluorophores, (2) the fluorescence emission couples into PC guided resonances and redirects the photons towards the collection objective for efficient extraction (3) the increased rate of spontaneous emission improves the rate of photon generation. This work also builds upon our recent advances with the design, synthesis, and application of Plasmonic Fluor (PF) nanoparticle tags, in which metallic nanorods are coated with a ∼3 nm thick dielectric spacer layer, followed by a coating of molecular fluorophores and target-specific capture molecules^27, 34, 35^. Here, we combine PCs and PFs as a hybrid system for the purpose of using PFs as digital-resolution tags in the context of a surface-based sandwich assay. We experimentally investigate the PC+PF hybrid system by constructing plasmonic nanostructures coated with a layer of fluorescent dyes and record the fluorescence enhancement on a PC substrate. Electromagnetic numerical simulations and a theoretical model of the hybrid enhancement are used to support experimentally measured enhancements, and to describe the design parameters for optimal coupling. We demonstrate the applicability of the method with the detection of human interluekine-6 (IL-6), a pro-inflammatory cytokine used in the diagnosis of cancer, sepsis, and autoimmune disease. We achieve a limit of detection of 10 fg/ml (representing approximately 20,000 molecules in a 100 μl assay volume) and a 7-log quantitative dynamic range. The PC+PF hybrid system provides a 52-fold signal enhancement compared to detecting PFs on an ordinary glass surface, which facilitates digital counting of PFs with a line-scanning instrument using an inexpensive 0.25 NA objective. The instrument scans a 1.2×1.0 mm^2^ sensor area in ∼5 minutes, thus overcoming the surface area limitations associated with oil-immersion lenses and resonators such as micro-rings. Our reported detection limit is three orders lower than conventional microplate ELISA and on a similar range to more sophisticated digital microbead approaches like Simoa^36^ for the same IL-6 analyte.

## RESULTS AND DISCUSSION

To exploit the photonic-plasmonic hybrid coupling towards enhancing the fluorescence of an emitter, we recognized key design criteria towards engineering the PC and PF nanostructures. It is critical for the PC band structure to be designed such that there are resonant leaky modes overlapping both the excitation and emission spectra of the chosen fluor^8, 9^. Furthermore, engineering constraints of utilizing an inexpensive lower NA objective (NA = 0.25) dictates the following criteria for the PC: (1) The existence of a resonant mode at the excitation wavelength, which can couple the input laser source at an angle lower than the NA of the objective. (2) Leaky resonant modes overlapping the fluor emission spectrum which out-couple the emitted fluorescence at angles under the angular collection bandwidth of the objective lens. (3) The PF resonance should encompass both the absorption and emission spectrum of the fluor^22^ to provide a combined advantage of enhanced near field excitation and an increase in the rate of emission through the Purcell effect.

Note that the three criteria require the PC and PF resonant frequencies (ω_*PC*_ *and* ω_*NC*_) to be strategically matched (ω_*PC*_ ≈ ω_*NC*_ = ω_*o*_). A further less obvious design criteria is: (4) Matching the radiative decay rate of the PC and the non-radiative decay rate of the PF. This can be more closely understood by describing the hybrid system using the temporal coupled mode theory (TCMT) ^26, 37^. Let us consider the hybrid coupling between the plasmonic nanostructure and the PC guided resonance (PCGR) with radiative (*γ*_*r*_) and non-radiative decay rates (*γ*_*nr*_). Since the PC is made of low-loss dielectric material, the non-radiative decay is assumed to be dominated by the lossy-metallic nanostructure (*γ*_*nr*_ ≈ *γ*_*ab*_). We can describe the amplitude of the oscillation (*a*) as:

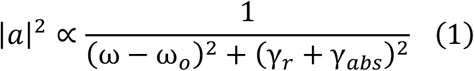

Using Eqn. 1, we can further evaluate the excitation enhancement at the resonant frequency (ω = ω_o_) ^38^ (details in Supplemental Note 1)

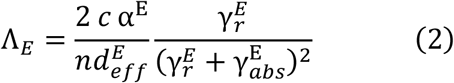

Where *n* is the refractive index of surrounding medium, 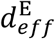 is the effective length of the evanescent field, and the α^E^ is the modal field confinement for the excitation mode. While evaluating enhanced extraction, the angular emission of molecules is altered due to the modification of the spectral density of states. Due to Helmholtz reciprocity, the outcoupled emitted photons from the resonator would be dictated by the mode solution obtained in Eqn. 1. By decomposing the Green’s function in the normalized Bloch mode basis with a finite resonance lifetime described by Eqn. 1, we can derive the enhanced rate of extraction (Λ_k_) for a specific (k, ω_*k*_) as (details in Supplemental Note 1):

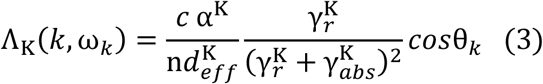

Where α^K^ is the energy confinement of the resonance mode and 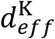 is the effective length of the evanescent field in the molecular layer in the extraction mode of emission. From Eqn. 2-3, we observe that the excitation (Λ_E_) and extraction (Λ_K_) enhancement are maximized when the rate of radiative decay in the PCGR mode is equal to the rate of non-radiative decay of the plasmonic nanostructure (*γ*_*r*_ = *γ*_*nr*_ = *γ*_*ab*_).

Based on this theoretical framework and design criteria, we engineered the two nanostructures individually and later integrated them to investigate the enhanced fluorescence. We first synthesized a stable fluorescently labelled PF. Following our third criteria, the PF is comprised of an inner gold nanorod core surrounded by a coating of silver, as shown in the transmission electron microscope (TEM) image in Fig. 2 (c). An additional siloxonane copolymer layer was used to create a spacer layer between the fluor layer and the metal surface. The optimal distance reduces the effect of metal induced fluorescence quenching while remaining in the vicinity of the strong near field. The LSPR of the synthesized particles was evaluated by measuring the extinction of the PFs on a TiO_2_-coated glass substrate to mirror the dielectric environment of the PC surface. As shown in Fig. 2(b), the PF exhibits a resonance near 660 nm, encompassing both the absorption and emission spectrum of the coated Cy5 dye.

**Figure 1.**
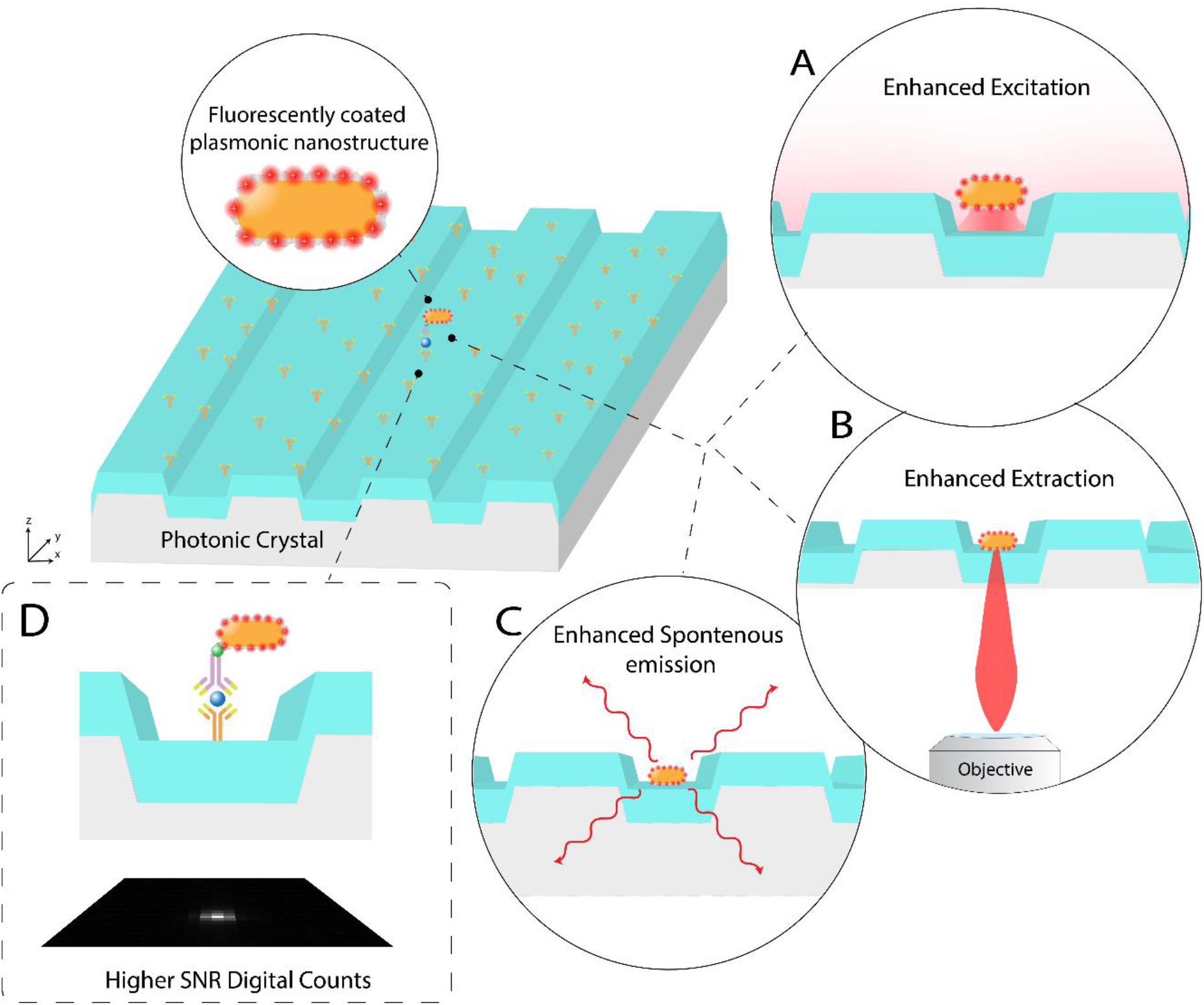
Principle of plasmonic-photonic hybrid for fluorescence enhancement applied towards a digital resolution immunoassay. (a) Enhanced excitation due to near field enhancement (b) Improved extraction efficiency towards a collection objective and (c) Increased rate of spontaneous emission. (d) The sandwich assay design utilized on the PC surface with PFs as fluorescent labelling tags for digital resolution detection.

**Figure 2.**
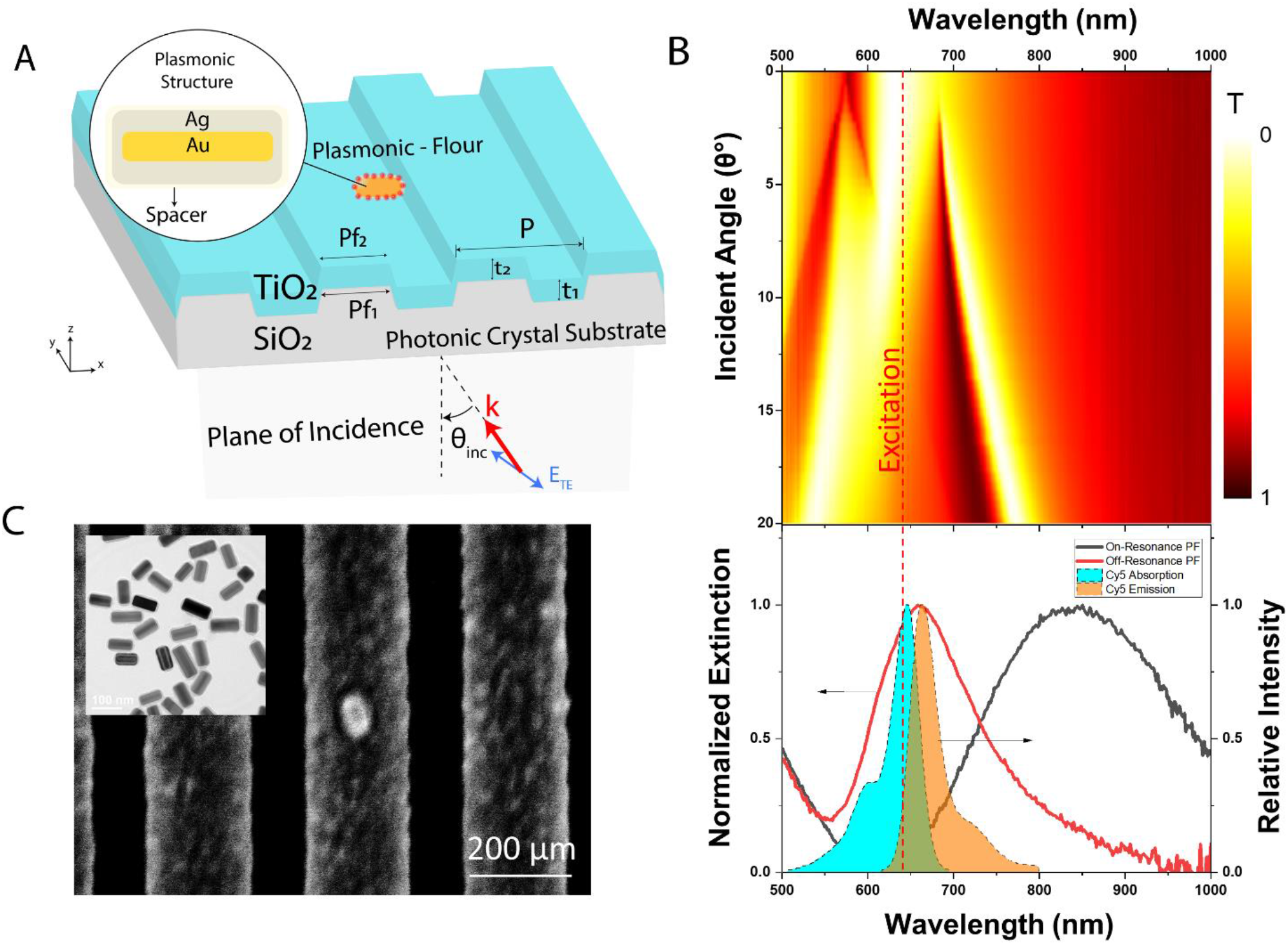
Design of the plasmonic-photonic hybrid for fluorescence enhancement. (a) PF on a PC surface. The structural parameters of the PC are: P = 380 nm, f_1_ = 0.34, f_2_ = 0.6, t_1_ = 80 nm, t_2_ = 114 nm. The excitation source is (λ_*ext*_ = 633 nm) is TE polarized E field at an incidence angle of θ_inc_ (b) Experimentally measured dispersion diagram from the transmission spectrum with the laser wavelength marked with a dashed red line. Lower panel: Extinction measurements of the on-resonance and off-resonance PFs with similar surface areas. The shaded regions correspond the absorption and emission spectra of Cy5 dye (c) Representative SEM image of a PF on the PC surface. Inset: TEM images of PFs.

Subsequently, we designed the PC following the requirement of using an inexpensive widefield 0.25 NA objective allowing a 15° window for excitation and extraction. The structure is composed of a periodically modulated glass substrate with a film of TiO_2_ (n = 2.4) deposited on top (Fig. 2 (a)(c)). The band structure was experimentally verified using far-field transmission measurements which shows close correspondence with the simulated band diagram (see Fig. S1). The PC exhibited a resonance at the excitation HeNe laser wavelength (λ_*ext*_) of 633 nm at near normal incidence for the transverse electric mode of excitation. Moreover, the radiative decay of this PC mode (*γ*_r_ = 7.08 × 10^14^ rads^−1^) was nearly equivalent to the estimated non-radiative decay rate (*γ*_ab_ = 7.44 × 10^14^ rads^−1^) of the NP (see Supplemental Note 1), achieving the optimal coupling condition previously derived from Eqn. 2.

A diluted concentration of streptavidin-coated PFs were conjugated on top of the PC surface coated with biotinylated-BSA (50 pg/ml) alongside additional BSA blocking to ensure a sparse density of nanoparticles (∼ 10^−3^/ μm^2^) and later dried. To experimentally interrogate the fluorescence enhancement resulting from different resonant modes of excitation, we built an epifluorescence detection instrument which provides precise control over the excitation angle in the *xz*-plane by translating the focused laser line along the back focal plane of a 10X (0.25 NA) objective lens (Fig. S2). The arrangement enabled us to couple the excitation laser through the phase matching condition into the guided resonance mode of the PC. The off-resonance condition was interrogated by tuning the angle of incidence away from the phase matching condition, effectively depriving the system of any guided resonance enhancement from the excitation source. Direct comparison to an unpatterned sample (incapable of enhanced excitation or directional extraction) was performed on a TiO_2_-coated glass substrate illuminated at normal incidence. A sample set of 100 individual PFs were imaged and processed to extract their peak fluorescence intensity for each mode of excitation, as shown in Fig. 3 (a).

**Figure 3.**
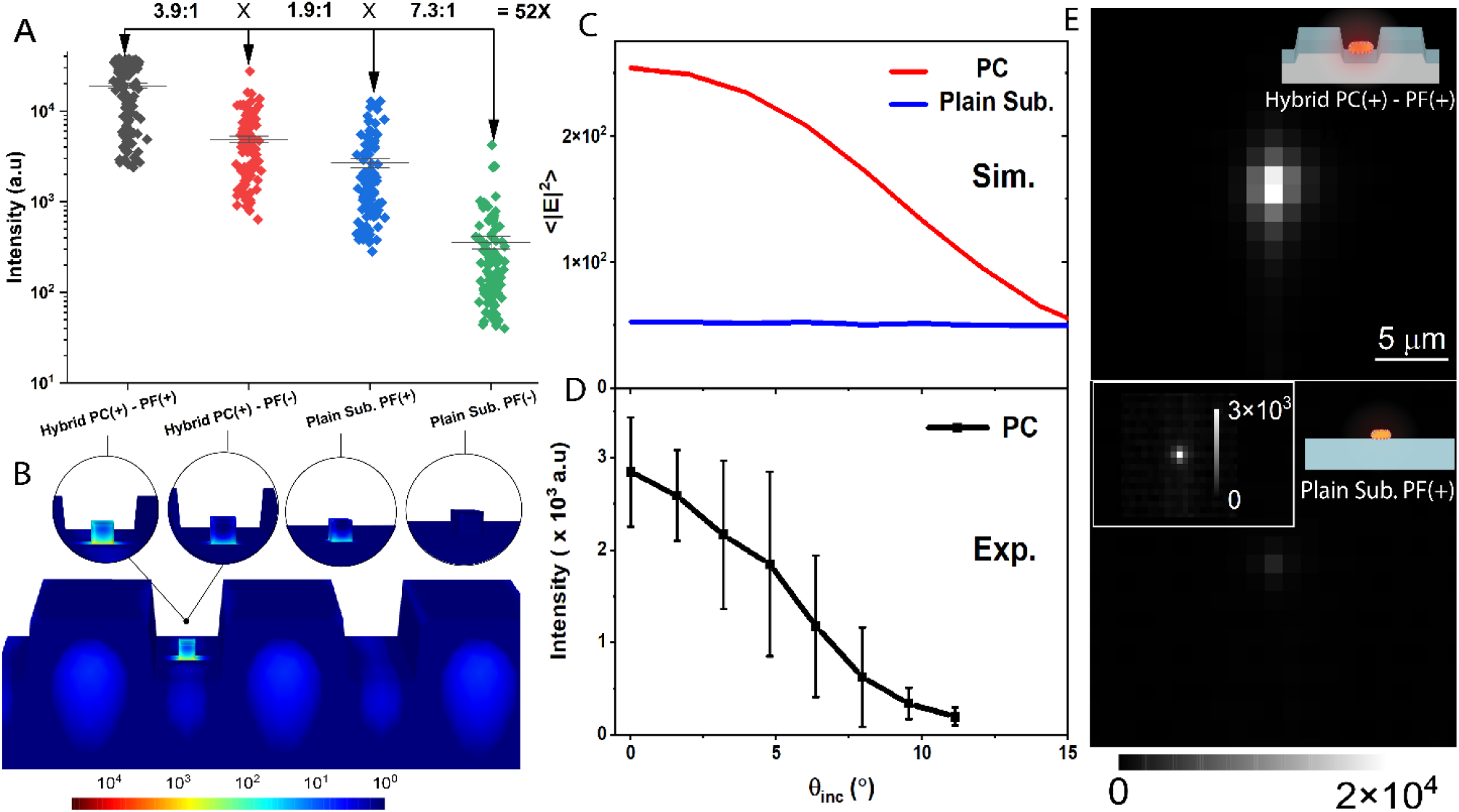
Fluorescence enhancement due to plasmonic-photonic coupling. (a) and (b) correspond to the experimentally measured peak fluorescence intensity and simulated near-field intensity (|E|^2^) for on-resonance (Hybrid PC(+) - PF(+) at θ_inc_ = 0°), off-resonance (Hybrid PC(-) - PF(+) at θ_inc_ = 13°) excitation of the PC and on-resonance (Plain Sub. PF(+) at θ_inc_ = 0°) and off-resonance PF (Plain Sub. PF(-) at θ_inc_ = 0°) on an unpatterned TiO_2_ coated glass substrate respectively. (c) The simulated <|E|^2^> near-field intensity for the excitation laser wavelength (λ_*ext*_ = 633 nm) at different angles of incidence. (d) Experimentally measured fluorescence intensity of the PF as a function of θ_inc_. (e) Representative fluorescence image of the PF when the on-resonance condition is engaged (top) and PF on an unpatterned TiO_2_ substrate (bottom).

We first sought to understand the effect of resonance matching of the plasmonic nanostructure with the absorption and emission spectra of the fluorescent dye. We employed off-resonant nanostructures with nearly the same surface area (σ_off_ = 1325 nm^2^) as the on-resonance particle (σ_off_ = 1206 nm^2^) which would ensure a similar density of fluorophore conjugation on both particles. The off-resonant particle exhibited a resonance at a wavelength of approximately 850 nm (Fig. 2 (b)), removing the possibility for surface plasmon resonance mediated enhancement. The peak fluorescence of both particles conjugated on an unpatterned substrate demonstrated 7-fold increase when the resonance condition is satisfied (Fig. 3 (a)). The enhancement is the result of both the increase in the excitation rate due to near field and the increase in the rate of spontaneous emission.

To investigate the contribution of Purcell enhancement, we carried out time-resolved photoluminescence measurements in the frequency domain. The average fluorescence lifetime for Cy5 and the on-resonance PF were 1 ns and 0.328 ns respectively (Fig. 4 (d)), resulting in a 2.8-fold increase in quantum efficiency, from 27% to 76% for the on-resonance PF (see Supplementary Note 3). The remainder of the enhancement can be attributed to the near-field enhancement due to the LSPR resonance. To evaluate the effect, full-wave electromagnetic simulations of the average density of near-field intensity inside the spacer layer < |E|^2^ > = ∫ E^2^ dr^3^/ ∫ dr^3^ showed a 10.9-fold increase when the plasmon resonance is closely matched to the fluor’s peak excitation wavelength (Fig. S4). Experimental measurements reveal a lower enhancement than the theoretically predicted value, which can be attributed to fluorescence quenching due its proximity with a lossy metallic surface. The field enhancement inside the spacer layer can be further amplified through the hybrid integration of the plasmonic nanostructure with a resonantly excited PC.

**Figure 4.**
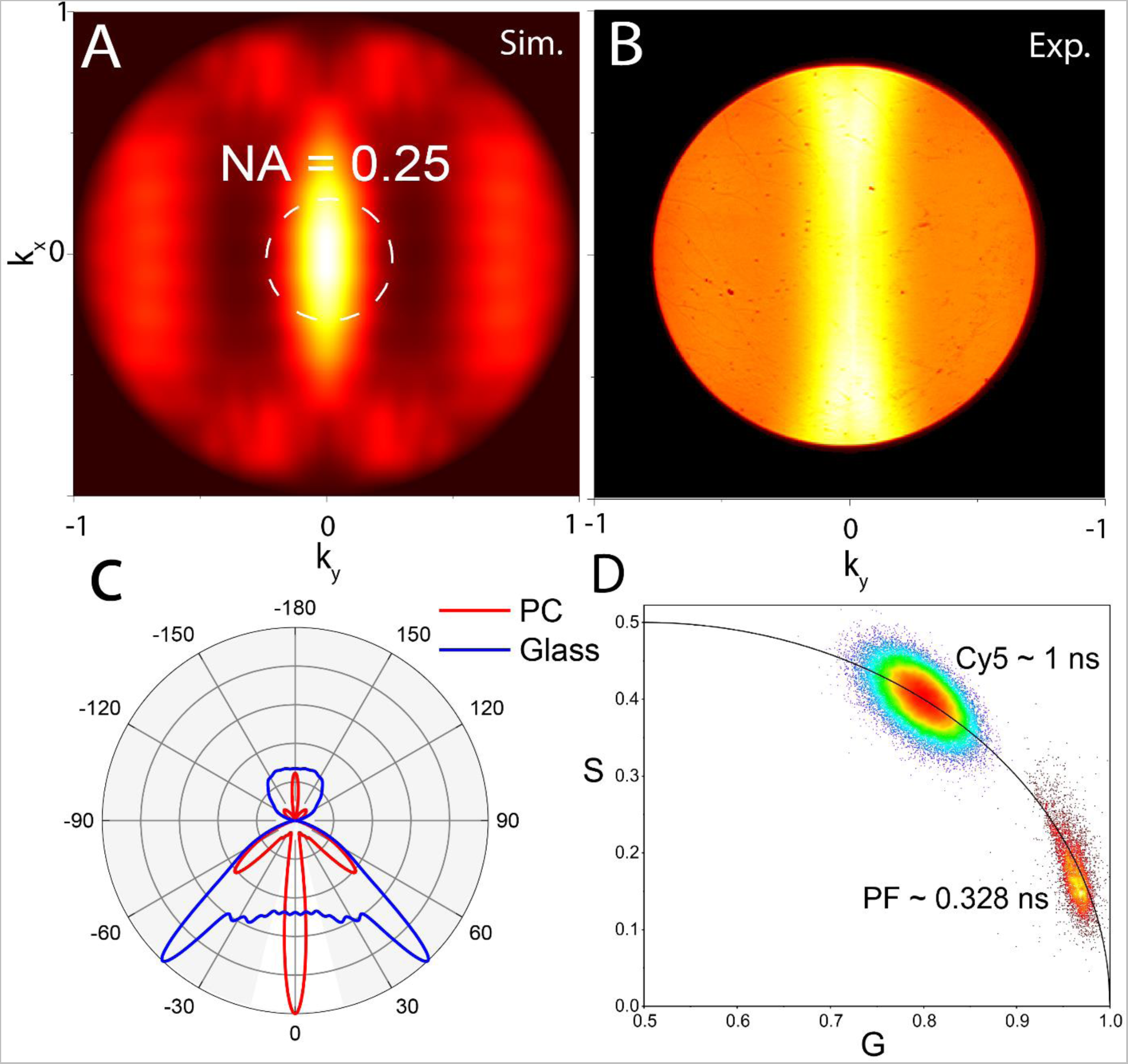
Enhancement due to extraction and the Purcell effect (a) Simulated Fourier plane image of PF on the PC surface (b) Experimentally measured back focal plane images of the PF in the TE mode of excitation (c) Simulated far-field radiation pattern of the PF on a PC and glass substrate. The white opening represents the allowed angular window for NA = 0.25 (d) Experimentally obtained phasor plot comparing the estimated lifetime of the Cy5 molecule and the PF.

The contributions of near-field enhancement due to the hybrid coupling can be understood by observing the angle-dependent resonance excitation of the PC. We observe that during the on-resonance excitation (θ_inc_ = 0°), the average fluorescence was 3.9-fold higher than the off-resonance (θ_inc_ = 13°) excitation (Fig. 3 (a)). This is the consequence of the PCGR mode coupling the excitation energy into a guided resonance and feeding the PF through a strong near-field interaction. The resonant coupling improves the efficiency with which the PF interacts with the input excitation. As a result, the surrounding spacer layer consisting of fluorescent dyes experiences an increased excitation energy. For further validation through numerical simulations, <|E|^2^> was evaluated for the hybrid coupled resonance, and a solitary nanoantenna on an unpatterned TiO_2_ substrate for the TE mode of excitation. The increase in the field intensity (λ_*ext*_ = 633 nm) around the nanorod can be observed when the phase matching condition with the incident wave excites the PCGR mode, whereas the value is angle-insensitive for the TiO_2_ substrate (Fig. 3 (c-d)). As a result, the PF tightly concentrates the field intensity with a maximum enhancement of over 10^4^ in certain regions of the spacer layer (Fig. 3 (b)). On comparison with the same nanostructure on an unpatterned TiO_2_ substrate, the simulated <|E|^2^> was 4.8-fold higher on the PC surface (Fig. 3 (c)), which is in good agreement with our experimentally obtained near-field enhancement (Fig. 3 (a)).

It is noteworthy that the broadened signal distribution observed in Fig. 3 (a) and Fig. 3 (d) can be attributed to two phenomena: (1) Variation of near-field intensity along the PC surface, resulting in a variability in the excitation energy depending on the relative position of the PF. (2) Polarization sensitive enhancement due to relative orientation of the PF with respect to the PC grating direction (see Fig. S5). One strategy to overcome these problems is the utilization of a 2D PC lattice design that could potentially provide a more uniform field distribution if two guided resonance modes are engineered to interact^39^. Moreover, the isotropic nature of a PC with 2D symmetry would eliminate the requirement of polarized excitation and reduce the effect of orientation-induced broadening of the intensities. Another strategy to minimize the effect of polarization sensitivity could be the utilization of a PF with more spherical symmetry, such as nano-cuboids that satisfy the same LSPR selection criteria as nanorods^40^.

We next investigated the far-field properties of the PC+PF hybrid system by characterizing the effects of directional emission extraction. On comparing the off-resonance fluorescence on the PC with the intensity measured from PFs on an unpatterned TiO_2_-coated glass substrate, we observed a 1.8X higher signal (Fig. 3 (a)), attributed to the coupling of the fluorescence emission with the leaky-PCGR modes which redirect the photons according to bandstructure as described in prior research^8, 16^. As a result, the angular emission for the dye follows the isofrequency contours of the PC, as described by the allowed resonance solution of the PC in momentum space (*k*_*x*_, *k*_*y*_) at a constant frequency ω.

In order to experimentally explore the k-space information of the emitted fluorescence, we imaged the back focal plane of the collection objective. We observe that the PC redirects the photons towards the normal axis of the collection objective, providing a more efficient pathway for collecting the emitted photons (Fig. 4 (b)). We further validated the experimental data with FDTD simulations to calculate the far-field radiation pattern and the collection efficiency (CE) for a PF on the PC. The modelling was performed by assuming a dipole source placed between the NP and the PC surface. The electric field was recorded below the lower grating and projected to the far-field to obtain (*E*(*k*_*x*_, *k*_*y*_)) The simulated data Fourier plane showed a good correlation with the experimental BFP (Fig. 4 (a-b)) and the 2D far-field radiation pattern (at *I*(*k*_*x*_ = 0, *k*_*y*_)) elucidates the difference between glass and PC surfaces (Fig. 4 (c)). The unpatterned surface has its major lobes near 45°, close to the critical angle of the glass/air interface. As a result, a large proportion of photons will be lost due to the reflection at this interface, resulting in poor collection efficiency for air-based objective. On the other hand, the PC band structure minimizes this effect by redirecting the photons towards the normal axis of the collection objective (Fig. 4 (c)). We further numerically calculated the CE as the ratio between collected power by the microscope objective (*S*_*col*_) and the total emitted power (*S*_*tot*_) by the fluorescent molecule 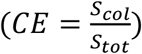. On comparing the two substrates at the peak fluorescence emission (λ_*em*_ = 665 nm), we estimate a 2X improvement in CE, correlating well with the experimentally obtained increase in fluorescence intensity (see details in Supplementary Note 3).

### Digital Resolution Bioassay

Of the numerous applications possible due to the hybrid enhancement, we demonstrate a digital resolution sandwich assay for the cytokine protein human Interleukin-6 (IL-6). IL-6 was selected for both its clinical utility as a diagnostic test for inflammation-related health conditions, and because it is a target biomolecule utilized in nearly all other detection technologies for performance benchmarking. The conventional enzyme/fluorescence-linked immunosorbent assay (ELISA/FLISA) utilizes a polystyrene microtiter plate with a capture antibody, biotinylated detection antibody, target analyte (Human IL-6) which is followed by exposure to an enzyme-substrate or streptavidin-fluorophore. In our method, the imaging substrate is replaced with a PC, while PFs are utilized as tags for the same assay design (Fig. 5 (a)). To efficiently conjugate capture antibodies, the TiO_x_ groups comprising the PC surface were chemically modified. Oxygen plasma activated PCs were silanized to achieve isocyanate functionality that facilitate antibody conjugation by reacting with the amine groups of lysine side chains. The corresponding urea linkage formed by this mechanism is stable at room temperature and resistant to further hydrolysis. Isocyanate groups that do not undergo rapid conjugation are efficiently converted to their corresponding amine via nucleophilic attack from surrounding water molecules, which promotes release of carbon dioxide. In order to maximize antibody immobilization density, we employed disuccinimidyl carbonate (DSC)^41^ as a secondary reaction component, which allowed for the regeneration of reactive isocyanates and succinimidyl carbamates that mutually facilitate immobilization via formation of a urea linkage between the antibody and the PC surface (see Fig. S6).

**Figure 5.**
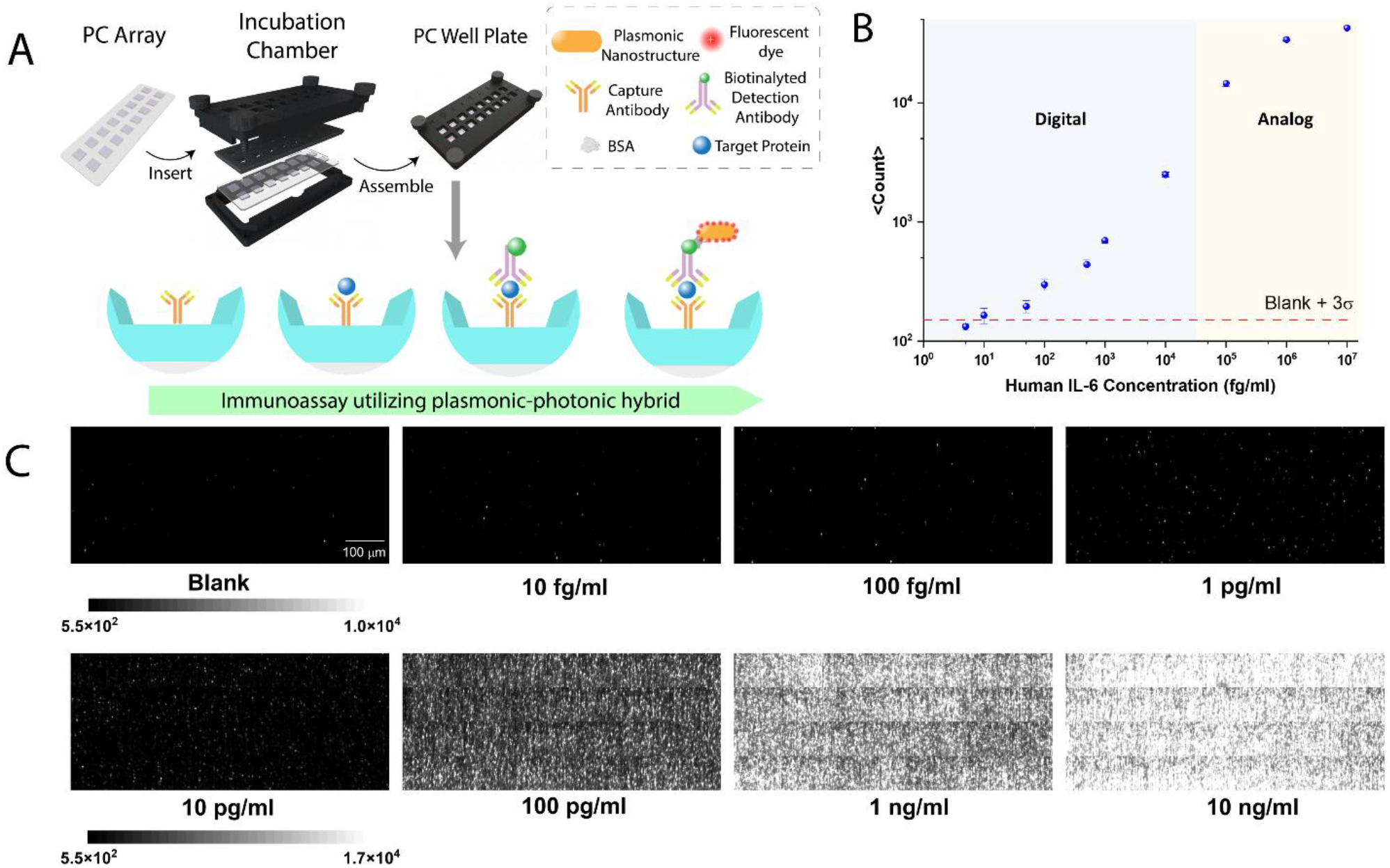
Digital resolution protein detection assay for IL-6 (a) Schematic illustration of the PF based immunoassay performed on a PC integrated within a multi-well plate. (b) Digital counts of the IL-6 immunoassay at different analyte concentrations. Blue region indicates the concentrations the algorithm utilized a digital counting method, and the orange area describes the region the analog intensity of fluorescent spot was used to estimate the digital count (3 repeats). (c) Representative fluorescence grey -scale images of a sub-area (0.4×1 mm^2^) showcasing the digital resolution PFs as a function of IL-6 concentration

To gauge the improvements in the sensitivity and the limit of detection, serial dilutions of IL-6 (10 ng ml^-1^ to 5 fg ml^-1^) were spiked into a PBS buffer solution to generate a dose-response standard curve. The assay was performed by incubating the 100 μl test sample upon the PC surface at room temperature for 2 hours, followed by a washing step with PBST buffer. A solution of 50 ng/ml biotinylated anti-IL6 were incubated on the PC surface for 2 hours, followed by a second washing step. Subsequently a solution of 2×10^7^ particles/ml streptavidin coated PFs were incubated for 30 minutes followed by a final washing step. A 1.2 × 1.0 mm^2^ area of the PC was scanned by tiling together 300 × 4.5 μm^2^ sub-images, and the appended images were processed to digitally count individual PFs (Fig. 5 (c)). Each sub-area scan required 100 msec, and the entire imaged area of 900 tiles required a scanning time of 5 minutes. The PFs are easily distinguished as individual units, where their signal-to-noise ratio (compared to the dark background of the non-fluorescent regions) is ∼55:1, highlighting the advantage of utilizing the PC+PF hybrid system to enhance the signal. We obtained a Limit of Detection (LOD) of 10 fg/ml, which was calculated by determining the concentration of target analyte with a digital count equal to 3 times the standard deviation above the mean of the blank control concentration (Fig. 5 (b)). The LOD represents six orders lower than the reported for conventional FLISA and three orders lower than reported for conventional ELISA^34^. We also note that the detection limit achieved is 500x lower than the 2 pg/ml LOD reported for the Luminex flow cytometer-based BeadArray™ technology^42^ and similar to the 4 fg/ml reported for the Quanterix Simoa™ technology^36^ that requires a much more complicated workflow that involves droplet partitioning, magnetic bead capture, and enzymatic amplification. Importantly, quantitation is maintained over a 7-log dynamic range, up to 10 ng/ml. Digital counting of PFs is utilized for the lowest (and most challenging) concentrations, up to a concentration of ∼10^4^ fg/ml. As expected, elevated concentrations result in a high density of captured IL-6 molecules, and thus a high density of PFs in the image. Thus, at elevated concentrations, the point spread functions of PFs in the image begin to overlap, and digital counting becomes less effective. However, quantitation can continue to higher concentrations by simply utilizing the “analog” average fluorescence intensity of the image, thus extending the dynamic range by a further 3-logs beyond the digital range. We note that this approach is much simpler and more direct than methods based upon making estimates of biomarker concentration using Poisson statistics and Average Enzymes per Bead (AEB) methodology^43^. It is interesting to note that the LOD of 10 fg/ml represents only ∼2.1 × 10^4^ molecules in the 100 μl assay volume, and that even with zero flow rate, our 120 minute stagnant incubation time measured 0.15% of the available molecules, assuming 100% efficiency for labeling the captured IL-6 molecules with a PF. Thus, we expect to achieve further reductions in LOD through liquid handling methods (such as stirring) that can help overcome diffusion-based molecular capture limitations.

A key advantage of the assay approach that the increased sensitivity does not entail any change in standard assay protocols. As a result, the method can be easily adopted toward a wide variety of biomarkers using sandwich-style assays (as shown here) for detection of proteins, or for detection of nucleic acid analytes (such as specific sequences of mRNA, miRNA, and ctDNA). Moreover, the platform is capable of digital resolution multiplexing in a single test region by utilizing PF tags that incorporate distinctly detectable fluorescence dyes for each analyte, while the PC can be prepared with an unpatterned mixed surface of capture molecules.

In summary, to overcome the limitation of weak fluorescence emission, we exploited the synergistic coupling between plasmonic and photonic crystal resonators to amplify the emission intensity of PFs by 52-fold. Through experimentation and numerical simulation, we attribute this increase to the strong near-field enhancement, improved extraction efficiency and boosted quantum yield. We harnessed the amplified intensity towards improving the signal-to-noise ratio of digital resolution immunoassays. By utilizing the plasmonic structures as a fluorescent reporter, integrated with photonic crystals as a readout platform, we obtained a limit of detection of 10 fg/ml for human IL-6. The signal enhancement elucidated here can be easily exploited to improve numerous existing bioanalytical techniques by enabling single-molecule level sensitivity while maintaining cost-effective instrumentation.

## METHODS

### PC Fabrication

The PC utilized in this work constituted a low refractive index grating coated with a higher refractive index (TiO_2_). The structure was fabricated on a glass wafer coated with a 10 nm etch stop layer of Al_2_O_3_. A subsequent SiO_2_ layer was deposited, and the periodic grating pattern was constructed by ultraviolet interference lithography carried out by Moxtek (Orem, USA). A TiO_2_ layer (thickness ∼ 114 nm) was then deposited on the etched wafer using sputtering to create guided waveguide structure.

### PF synthesis

Silver cuboids (AuNRAg) (synthesis in Supplement Note 2) were employed as the plasmonic core to prepare PF. 1 μl of (3-mercaptopropyl)trimethoxysilane (MPTMS) (Sigma Aldrich, 175617) and 1 ml of AuNRAg (extinction ∼2) were mixed and shaken on rocking bed for 1 hour. Next, 2 μl of APTMS (Sigma Aldrich, 281778) and 2 μl of trimethoxy(propyl)silane (TMPS) (Sigma Aldrich, 662275) was added to the MPTMS-modified AuNRAg to form the polymer spacer layer. Excess monomers were removed from the AuNRAg–polymer solution by two centrifugations at 6000 rpm for 10 min. After each wash the pellet was redispersed in 1 M CTAC aqueous solution to ensure colloidal stability. Polymer coated AuNRAgs were concentrated into a final volume of 5 μl. Next, to conjugate Cy5-BSA-Biotin complex (synthesis in Supplementary Note 2) to polymer modified AuNRAgs, we followed procedures mentioned in previously reported study^44^. Briefly, to allow coating of Cy5-BSA-Biotin to AuNRAgs, the pH of 100 μl 4 mg ml^−1^ Cy5-BSA-Biotin was lowered by adding 1 μl of 20 mg ml^−1^ citric acid (Alfa Aesar, 36664). To this solution, concentrated AuNRAg-polymer solution was added, and the resulting solution was sonicated for 20 min in dark. After coating, excess Cy5-BSA-Biotin was removed by centrifugation at 3,000 rpm for 10 min and incubated with 0.4 mg ml^−1^ of Cy5-BSA-Biotin in pH 10 nanopure water (1 μl NaOH in 10 ml of water) for 3 days at 4 °C. Finally, the nanostructures were washed 4 times using pH 10 nanopure water by centrifugation at 3,000 rpm for 10 mins. The nanolabels were then redispersed in 1% BSA in 1X PBS solution for use in immunoassays. To further label streptavidin onto these nanolabels, the biotinylated plasmonic-fluor solution was incubated in 100 μg/ml solution of streptavidin for an hour on the rocking bad and subsequently purified by washing 4 times using pH 10 nanopure water by centrifugation at 3,000 rpm for 10 mins.

### Spectroscopic measurements

For the PC transmission band diagram, the sample was mounted on a fine-resolution motorized rotation stage. A deuterium and halogen lamp (Ocean Insight DH-2000-BAL) was used to produce a white light collimated beam of 5 mm^2^ area with TE polarization. The incident angle was tuned with computer controlled motorized rotation stage between θ_*i*nc_ = 0°-14°. The zero-order transmission was collected by a fiber collimating lens and guided to a spectrometer (Ocean Optics USB 2000) to measure the spectral intensity.

The LSPR measurements of the PF samples were carried out by drop casting them on a TiO_2_ coated glass slide to mimic the dielectric environment of the PC surface. The extinction spectrum mas measured using a micro spectrometer setup built on a Zeiss Axio Obsever D1 inverted microscope. A sparse density region of the sample was identified, and the transmitted intensity was collected 20X objective lens and directed to a silicon PDA spectrometer.

### Numerical simulations

To simulate the near-field properties, we carried out finite-element-method simulations (COMSOL Multiphysics). The unit cell of the simulation spanned three PC periods (1.14 um) in the *x*-direction and 800 nm in the *y*-direction. The geometric parameters were inferred from atomic force microscope (AFM) measurements and by further best-fitting with the far-field transmission spectra of the bare PC. The refractive index of TiO_2_ (n = 2.44) was taken from Seifke^45^ and the SiO_2_ (n = 1.44) were referred from the manufacturer. The plasmonic nanostructures were extracted from the TEM images and the optical constants of gold and silver taken from Johnson and Christy. <|E|^2^> was evaluated in a 2 nm spacer layer with a refractive index of 1.5. The full field solutions of the PC were first calculated assuming Floquet periodic boundary conditions. The field distribution was subsequently used as a background field for the nanoparticle excitation with the Floquet boundary condition replaced with perfectly matched layers.

The far-field properties were simulated using finite difference time domain method (Lumerical FDTD). The Fourier plane field distribution was calculated by assuming a monochromatic dipole source placed in a 2 nm gap between the plasmonic flour and the photonic crystal surface. The unit cell consisted of 50 periods, modelling the coupled emission to traverse along the waveguide for a finite lifetime. The structure was surrounded with perfectly matched layers and electric field above and below the plane of the PC were recorded. The field profile was projected from the near-field to the far-field to obtain the k-space intensity (*I*(*k*_*x*_, *k*_*y*_, *λ*)) information and was further averaged over the fluorescence emission spectrum of the Cy5 to obtain *I*(*k*_*x*_, *k*_*y*_). The collection efficiency was further calculated as a ratio of the integrated power per unit solid angle across the angular bandwidth of the collection objective and the total solid angle. Furthermore, to account for the losses encountered by the glass/air interface, the Fresnel coefficient was applied to the field distribution (See supplementary note 3).

### Imaging setup

The setup was custom built line-focused epifluorescence microscope setup with a 10X objective (0.25 NA, Olympus LMPLFLN) used for both the excitation and collecting emitted light (see Fig. S2). A red excitation HeNe laser (λ_*exc*_= 633 nm) was beam expanded to ∼1 cm by a two-lens relay. It was subsequently linearly polarized to the transverse electric mode and a half-waveplate provided control over the illumination power. A cylindrical lens focused the beam to the back focal plane of the objective lens which resulted in a focused beam along *xz*-plane and collimated excitation along the *yz-*plane of the sample. A motorized translational stage allowed us to tune the angle of excitation by shifting the position of the focused line beam on the back focal plane. The emitted fluorescence was collection by the objective and an emission notch filter was used to reduce the background laser excitation. A tube lens focused the image plane onto a CCD camera (Synapse EMCCD) which recorded the images. The laser was focused to a line of the area 300 μm x 4.545 μm area. A series of images were acquired by translating sample in the *xy*-plane in steps of the focused area resulting in an appended field of view of 1.2 mm x 1 mm.

### Image analysis

The raw images were first Wiener filtered to reduce the non-uniformity appending created by the line scanning. The maximally stable extremal regions (MSER) were adopted for the blob detection algorithm which were binarized above the noise levels of the image. The morphological property and size were used as a selection criterion to determine the count and gauge on the peak, total and average intensity values (see Fig. S3). The signal-to-noise comparisons in the fluorescence enhancement were done by subtracting the background noise intensity of the unconjugated plain substrates with the focused laser 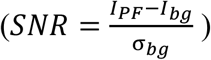.

### Lifetime measurements

The lifetime data in this study were measured using digital frequency domain fluorescence lifetime imaging microscope (FastFLIM, ISS) and analysis with phasor plots. The time-resolved photoluminescence (TRPL) was measured from single PFs or homogeneously distributed dye layer. Using Digital Frequency Domain (DFD) technique allows for the acquisition of Time-Tagged-Time-Resolved (TTTR) data without the dead time typical of Time-correlated single-photon counting (TCSPC) approach. The excitation source was a pulsed single photon laser with single-mode fiber optic output and optical parametric oscillator, producing 100-120 fs pulses at λ_laser_ = 635 nm and a repetition rate of 80 MHz. The emission was collected a high-NA objective lens (NA=1.46, oil immersion, 100X, Zeiss α Plan-APO-CHROMAT) and passed through a band-pass filter (692nm CWL, 40nm Bandwidth) to remove the excitation laser and imaged onto silicon single-photon avalanche photodiodes (Si-SP-APDs). 692nm CWL, 25mm Dia, 40nm Bandwidth. The Single-photon avalanche diodes (SPADs) were connected to a photon counting module to acquire the Time-Tagged-Time-Resolved (TTTR) data to generate the histogram of photon arrival times. The raw FLIM measurements from each pixel are directly located on a 2D phasor plot (Figure 4 (d)). This relationship is represented as a semicircle curve centering at (G=0.5, S=0) with a radius of 0.5 on the phasor plot. The semicircle curve indicates the lifetime trajectory with decreasing lifetime from left to right, where (1, 0) indicates lifetimes near zero to (0, 0) being infinite lifetime.

### Surface functionalization

A Human IL-6 DuoSet ELISA kit (R&D, catalogue number DY206, lot P173353) was used to test PF capture on PCs. The PCs were diced into 3 by 4 mm chips and glued onto coverslips using Norland Optical Adhesive 63 (Thorlabs), with UV curing for 3 minutes. The PC surface was first washed by sonication in acetone, isopropyl alcohol, and MilliQ water for two minutes each, then dried at 120°C for ten minutes. The surface was oxygen plasma-treated for 10 minutes at 100% power in a PicoDiener system, and silanized with a 5% silane mixture (95:5 ratio of 3-(triethoxysilyl)propyl-isocyanate to chlorobutyldimethyl silane, from Millipore Sigma in tetrahydrofuran for 30 minutes at room temperature. The surface was washed again by sonication in acetone, ethanol, and MilliQ water, and wells were added using ProChamber Mircoarray system.

### Sandwich ELISA for Human IL-6 Detection

Capture antibodies were added from the DuoSet kit at a concentration of 2 μg/mL in 1xPBS, 100 nM N,N’-disuccinimidyl carbonate for 2 hours at room temperature. The surface was washed five times with PBST, followed by blocking with 200 μl of reagent diluent (1xPBS containing 3% BSA, filtered by 0.2 μm). Washing five times with PBST was repeated, and 100 μl of serially diluted standard solution (tenfold dilution from 1 fg/ml to 10 ng/ml using reagent diluent) were added into the wells and incubated for 2 hours at room temperature. Washing with PBST was repeated after standard human IL-6 incubation, then 200 μl of biotinylated detection antibodies (50 ng/ml in reagent diluent) were incubated for 2 hours. The PCs were washed again and incubated with a 2×10^7^ particles/ml concentration of Cy5-plasmonic fluors in reagent diluent for 30 minutes. A final washing step is repeated, then 200 μl of 1xPBS were added to each well until imaging.

## ASSOCIATED CONTENT

### Supplementary information

The supporting information file is available free of charge – Supplemental_Information.pdf Supplementary information includes – Simulation of PC band diagram; schematic of optical setup; image processing and counting algorithm; simulation of the plasmon resonance; variation of the electric field density along the grating; surface functionalization chemistry; derivation of the theoretical framework; detailed synthesis of the PF; collection efficiency and quantum yield calculation; optimization steps for surface conjugation of antibodies.

## AUTHOR INFORMATION

### Notes

The authors declare the following competing financial interest(s): S. Singamaneni is an inventor on a pending patent related to plasmonic-fluor technology and the technology has been licensed by the Office of Technology Management at Washington University in St. Louis to Auragent Bioscience LLC. S. Singamaneni is a co-founder/shareholder of Auragent Bioscience LLC. S. Singamaneni along with Washington University may have financial gain through Auragent Bioscience, LLC through this licensing agreement. These potential conflicts of interest have been disclosed and are being managed by Washington University in St. Louis.

### Author Contributions

B.T.C., S. Singamaneni, Y.X. and P.B. designed the study. P.B. and Y.X. carried out the optical characterization experiments for the enhanced fluorescence. P.B. performed the numerical simulations and outlined the theory with assistance from Y.X.. S. Shepherd. and P.B. optimized the surface chemistry. S. Shepherd., P.B., R.G., and Y.X. ran the protein detection experiments. R.G. synthesized and characterized the plasmonic-flour particles. Y.X. conducted the BFP enhanced extraction experiments and Purcell factor experiments. L.A., J.T., Y.X., and H.K.L. helped in key steps while designing the PC well plate and bioassay experiments. P.B. drafted the manuscript with the assistance of all the authors.

## ACKNOWLEDGEMENT

The work was supported by National Science Foundation (1900277), National Institute of Health (NIH R01 5R01CA227699-03), and Cancer Center at Illinois (CCIL). Y.X is grateful for the C*STAR fellowship from CCIL and J.T acknowledges support from the NSF graduate fellowship program. The authors also acknowledge Nantao Li, Congnyu Che, Glenn. A. Fried, Leyang Liu, Pin Ren, Shengyan Liu, Hanwei Wang, and Qinglan Huang at the University of Illinois at Urbana Champaign and Yuansheng Sun at the ISS inc. for their valuable discussions.

