## Supplementary Information for "Photonic-Plasmonic Coupling Enhanced Fluorescence Enabling Digital-Resolution Ultrasensitive Protein Detection"

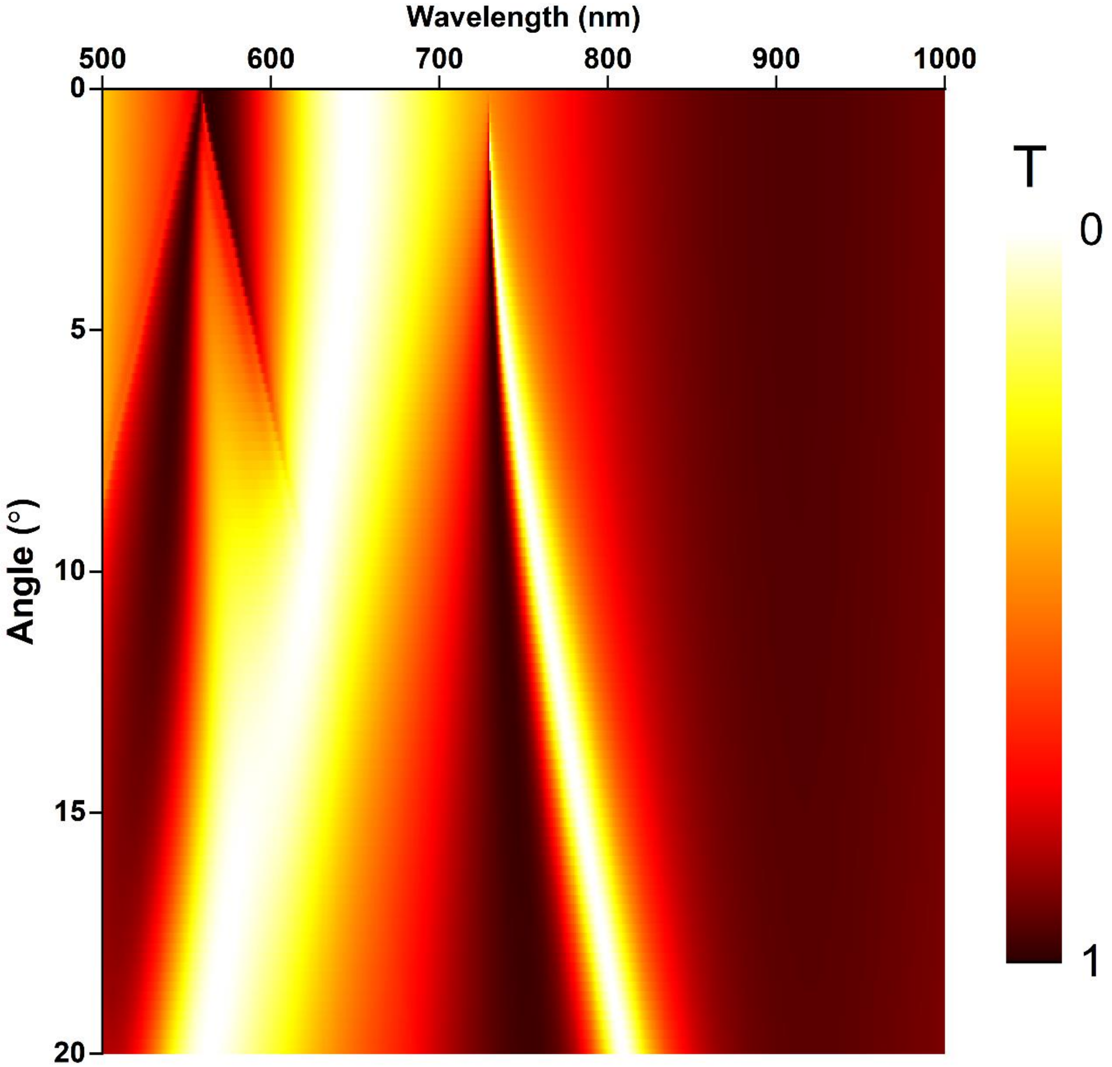


**Figure S1**. Numerical simulation of the PC band diagram using the structural properties shown in Fig. 2 in the TE mode excitation performed using COMSOL FEM simulations.

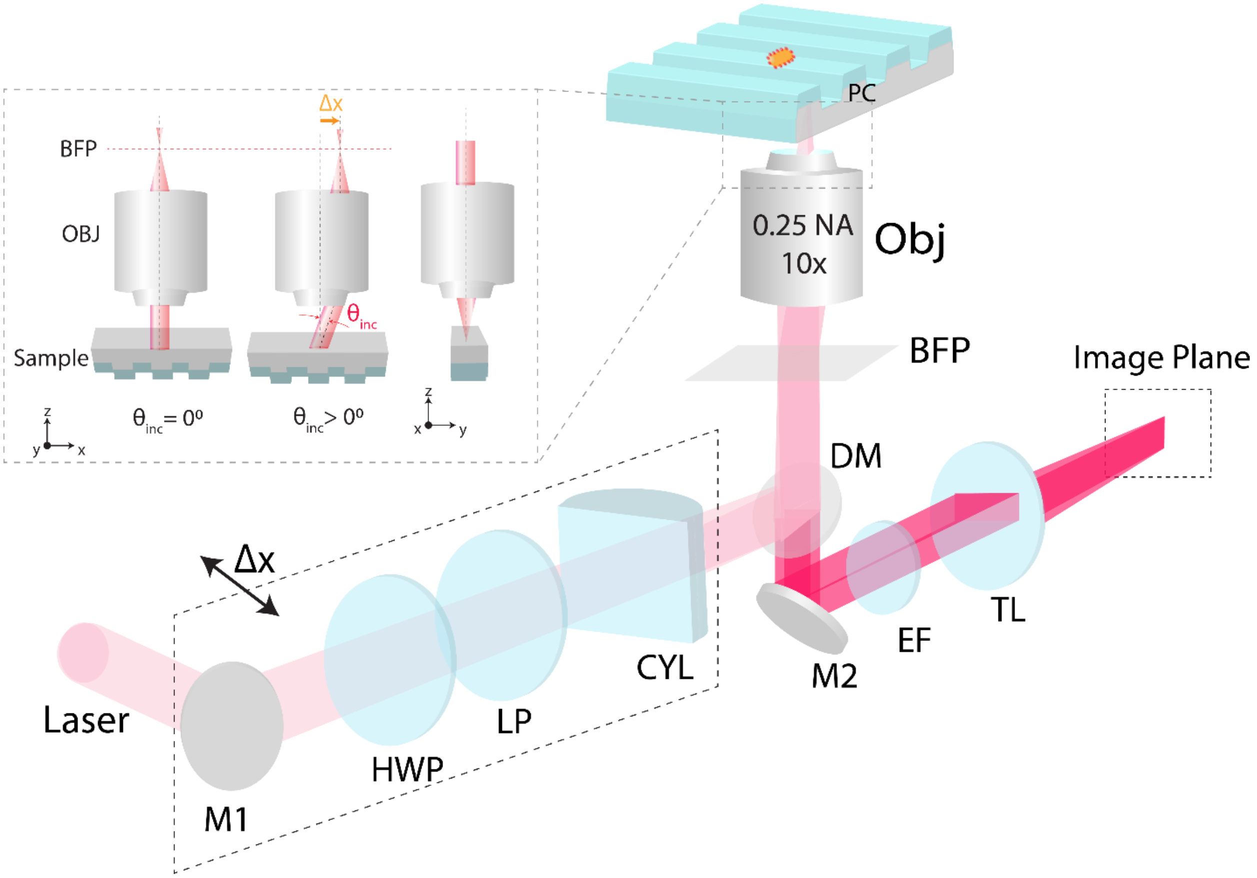


**Figure S2.** Schematic diagram optical setup used to investigate the plasmonic-photonic coupling. Where the M: Mirror, LP: Linear Polarizer, CYL: Cylindrical lens, EF: Emission Filter, TL: Tube lens, Obj: Objective. The optical components described in box are mounted on a motion stage which can translated along the x- axis. The inset showcases the angle tuning mechanism by spatially moving the focused light in the BFP of the objective lens.

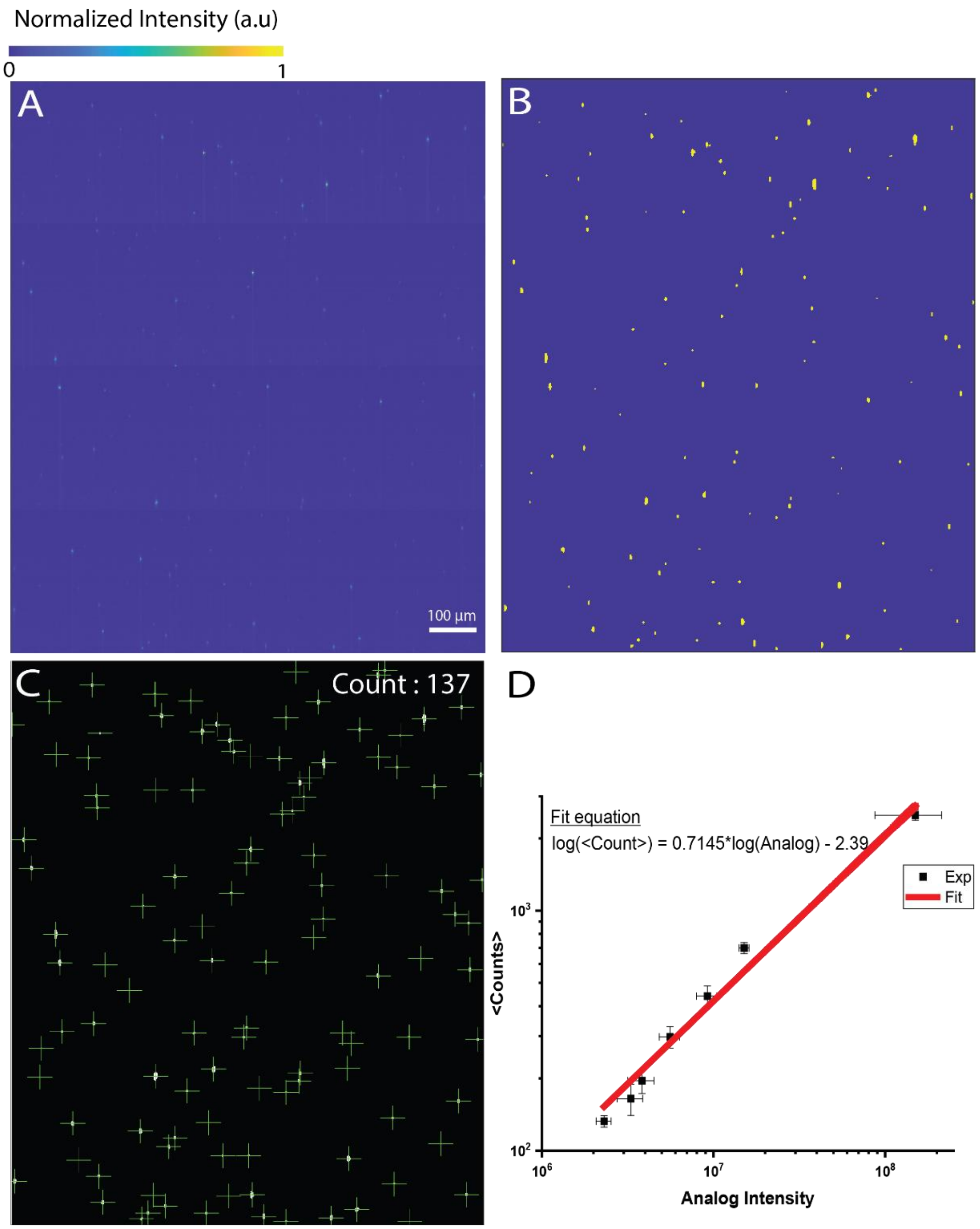


**Figure S3.** Image processing and counting algorithm of obtained fluorescence intensity (a) shows the original fluorescence image of individual PFs on the PC substrate (b) the blob detection algorithm binarizing the image for the pixels about a certain threshold (c) The cross marks indicate the centroid of the detected particles with the estimated counts described on the top right (d) Equation utilized in estimating the counts for higher concentrations (>$10^5$ pg/ml) based on the log-log linear relation obtained between the analog and digital counts for the lower concentrations (<$10^5$ pg/ml). The analog intensity for the lower concentrations was calculated using the sum of the fluorescence intensities of all the individual binarized blobs as described in Fig. S3 (b).

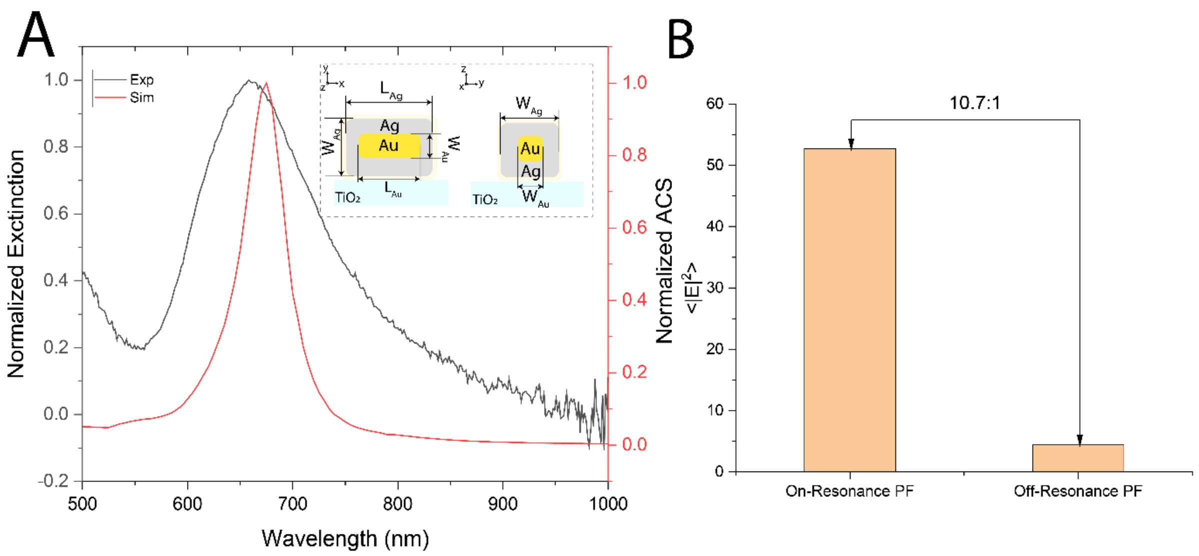


**Figure S4**. The experimentally measured and simulated resonance position of the plasmonic nanostructure used in the analysis with $W_{Ag}$ = 30 nm, $W_{Au}$ = 10 nm , $L_{Au}$ = 60 nm, $L_{Ag}$ = 70 nm, Spacer layer thickness = 2 nm. (b) Simulated increase in the average near field intensity inside the spacer layer (<|E|$^2$>) while comparing the on-resonance PF ($L_{Ag}$ = 70 nm) and the off-resonance PF ($L_{Ag}$ = 100 nm).

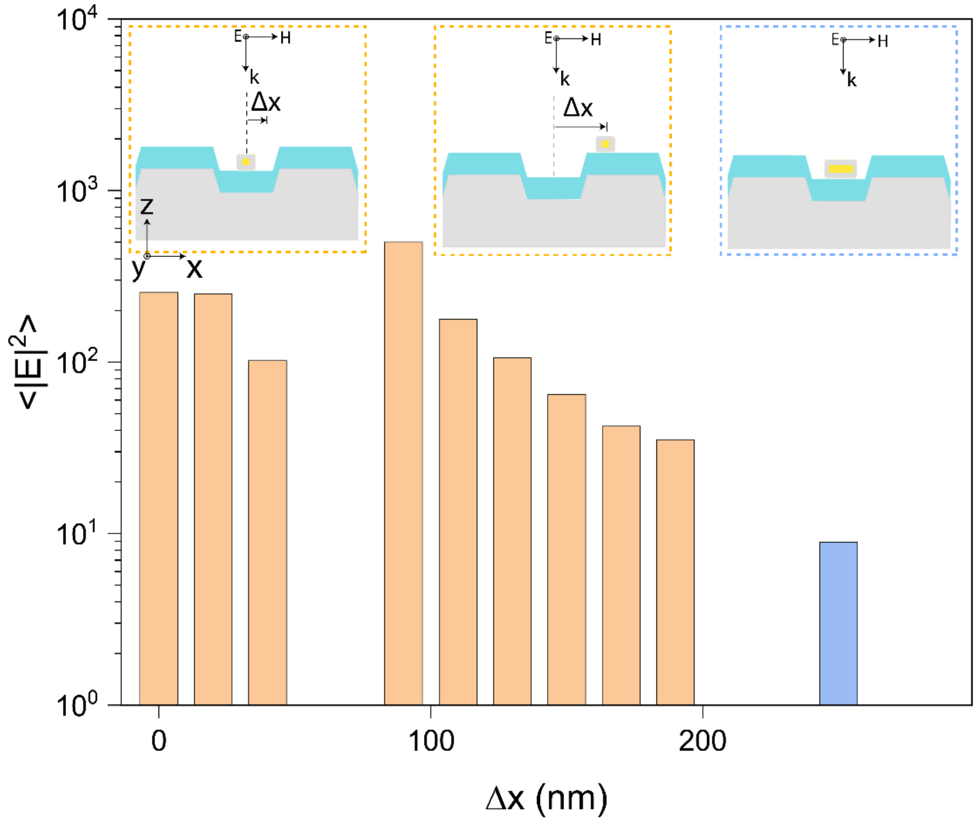


**Figure S5**. The simulated average electric density inside the spacer layer as the plasmonic nanostructure is moved along the grating structure of the PC. The orange and blue bar indicate the case when NP is aligned along and perpendicular to the direction of excitation electric field respectively

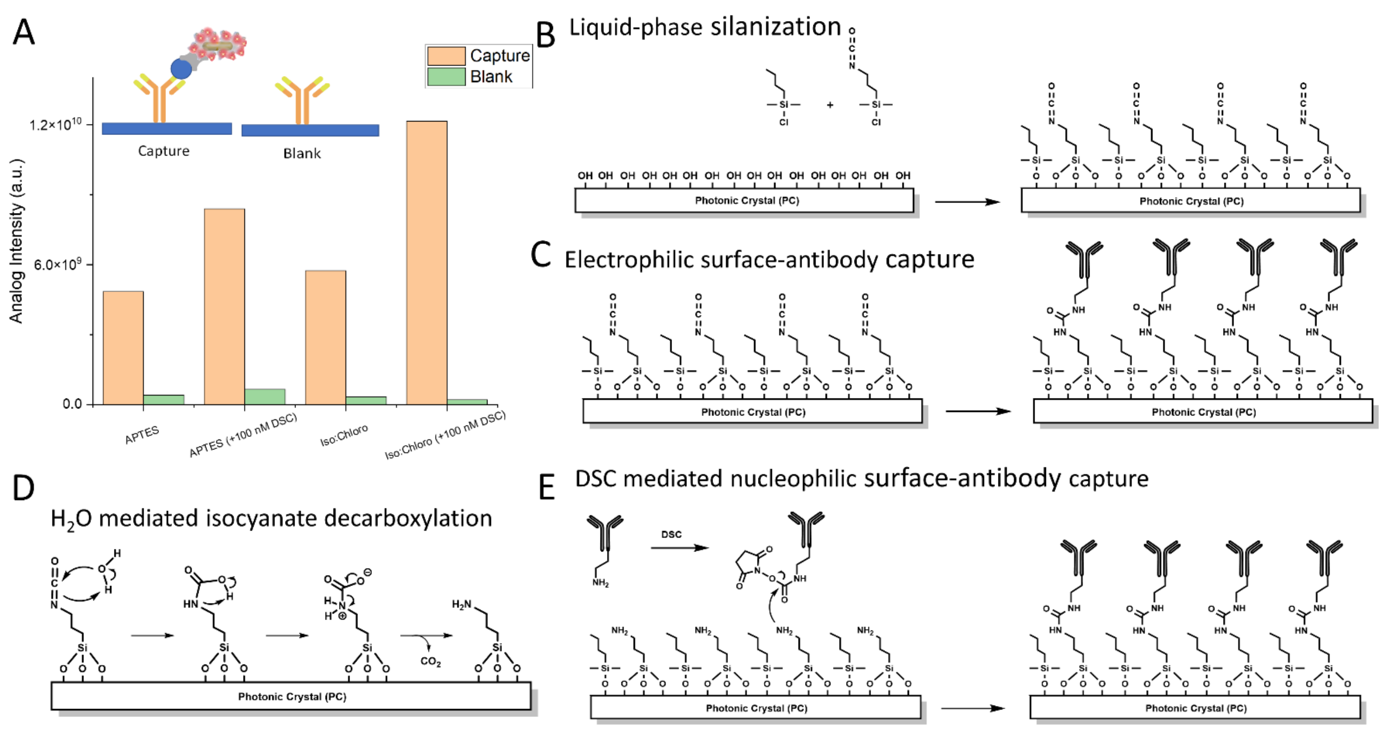


**Figure S6.** Surface functionalization chemistry (a) shows the variation in surface density of capture antibodies on the PC for different surface chemistries (b) describes the liquid phase salinization carried out on plasma treated PC surface using (3-Isocyanatopropyl)triethoxysilane (Iso) and butyl(chloro)dimethylsilane (Chloro) (c) the electrophilic surface-capture of the antibody with the isocyanate functional group (d) the mechanism of the $H_2O$ mediated isocyanate decarboxylation (e) Nucleophilic surface-capture of the antibody mediated through the DSC linker.

**Supplementary Note 1: Theoretical framework**

<u>Excitation</u>

In this section, we elucidate the theoretical understanding of the hybrid system for excitation enhancement using the temporal coupled mode theory [1-3]. Considering a PC cavity and AuNR with a resonant frequency $\omega_{PC} = \omega_{NC} = \omega_o$. The PC having a radiative decay rate of $\gamma_r$ and the nanoantenna with a non-radiative decay rate ($\gamma_r$ = $\gamma_{abs}$). By modelling the hybrid resonance as a resonator which exchanges energy with an incoming wave ($s_{1+}$) with coupling constants $\kappa$ we can obtain

$$\frac{d}{dt}a = (-i\omega_o - \gamma_r - \gamma_{abs})a + \kappa s_{1+} \qquad (1)$$

Where a is the amplitude of oscillation. Considering only a single input excitation $s_{1+} = s_{1+}e^{-i\omega t}$ at frequency ω

$$|\mathrm{s}_{1+}|^2 = \frac{1}{2}\epsilon_o|\mathrm{E}_{\mathrm{inc}}|^2 \mathrm{cA_i} \qquad (2)$$

Where the power of the normal incident field is $E_{inc}$ and the area of cross-section $A_i$. In Eqn. 1, the coupling coefficient with the excitation channel can be given as $\kappa = \sqrt{\gamma_i}$, where the $\gamma_i$ is the contribution of the total radiative decay by the excitation channel. Under a first approximation consider a symmetric two-sided cavity, one can estimate[4] $\kappa = \gamma_i^2 = \sqrt{(\gamma_r/2)(A_c/A_i)}$. On evaluating the equations in the steady state, the stored energy in the resonators

$$|\mathrm{a}|^2 = \frac{1}{(\omega - \omega_o)^2 + (\gamma_r + \gamma_{abs})^2}\kappa_1^2|\mathrm{s}_{1+}|^2 = \frac{1}{2}\epsilon_o|\mathrm{E}_{\mathrm{loc}}|^2 \mathrm{V_{eff}} \qquad (3)$$

Where the $E_{loc}$ is the maximum local field amplitude and $V_{eff}$ is the effective modal volume. Using Eqn. 3, we can obtain the field enhancement on resonance[3]

$$\Lambda_E = \frac{|E_{loc}|^2}{|E_{inc}|^2} = \frac{2A_c}{\pi V_{eff}^E}\frac{\gamma_r^E}{(\gamma_r^E + \gamma_{abs}^E)^2} = \frac{2c\alpha^E}{nd_{eff}^E}\frac{\gamma_r^E}{(\gamma_r^E + \gamma_{abs}^E)^2} \qquad (4)$$

Where $\alpha^E = \int |E^E(r)|^2 dr$ is the modal energy confinement and $d_{eff}$ is the length of the effective evanescent tail. The decay rates for the PC and the nanoantenna were derived experimentally determining the full width half maxima of their individual resonance spectrums $\gamma = \frac{\Delta\omega}{2}$. For the PC at normal angle of incidence with resonance centered near the 633 nm excitation (Fig. 2 (b)), the FWHM ($\Delta f = 90$ THz) results in a $\gamma_r = 7.08 \times 10^{14}$ rads$^{-1}$. The FWHM ($\Delta f = 94.7$ THz) for the plasmonic structure was calculated from the extinction spectrum (Fig. 2 (b)) resulted in the $\gamma_{abs} = 7.44 \times 10^{14}$ rads$^{-1}$.

Extraction

To understand the effect of extraction, we consider the modification of spectral density of states due to the presence of the photonic crystal. On coupling with a resonance, the rate of photons generated by an isotropic collection of molecules can be derived by applying the Fermi's golden rule in the normalized bloch modes[5]. The decay rate $\Gamma^{PC}$ for an ensemble of molecules $N_o$ can be written as:

$$\Gamma^{PC} = N_o \int E^2(r) \Delta N \, dr \quad (5)$$

Where the $\Delta N$ is the number of leaky guided modes of the PC at particular guided mode vector. On evaluating Eqn. 5 considering the number of modes in a given linewidth $\Delta\omega$ with density of states taking a Lorentzian line shape

$$\Gamma^{PC}(\omega) = \frac{N_o A \omega |\mu|^2}{6\pi^2 \hbar \epsilon_o} \int_k \left( \int |E(r)|^2 dr \frac{\Delta\omega_k}{(\omega - \omega_k)^2 + \gamma^2} \right) dk \quad (6)$$

Where μ is the moment dipole, A is the active area. On evaluating Eqn. (6) at the resonance of the PC[3] ($\omega = \omega_k$)

$$\Gamma^{PC}(k, \omega_k) = \frac{N_o A \omega N_o |\mu|^2}{6\pi^2 \hbar \epsilon} \frac{\alpha^K}{\gamma^K} cos\theta_k \quad (7)$$

$\alpha^K = \int |E^K(r)|^2 dr$ is the energy confinement in the gain medium for the fluorescence resonance mode, and $\theta_k$ is the angle made at the particular k. It is noteworthy, only a fraction of generated photons would coherently radiated to the far-field due to the non-radiative decay in the plasmonic nanoantenna which can be accounted as $\Gamma^{PC} \times \frac{\gamma_r^K}{\gamma^K}$. The overall enhancement of the extraction rate, for a particular k, in presence of the PC compared to the free space[3, 5]

$$\Lambda_K = \frac{\Gamma^{PC} \times \frac{\gamma_r^K}{\gamma^K}}{\Gamma^{free}} = \frac{c\alpha^K}{nd_{eff}^K} \frac{\gamma_r^K}{(\gamma_r^K + \gamma_{abs}^K)^2} \cos\theta_k$$

**Supplementary Note 2: Detailed synthesis process of PF**

Synthesis of plasmonic core:

To prepare plasmonic core specific to Cy5 dye employed in the study, first AuNRs with absorbance wavelength 950 nm (localized surface-plasmon resonance wavelength) were synthesized by the seed-mediated method[6]. If needed, the wavelength of gold nanorods can be easily tuned to couple with different dye molecules to achieve the best enhancement factor. Briefly, to a solution of 9.75 ml 0.1 M hexadecyltrimethylammonium bromide (CTAB) (Sigma Aldrich, H5882), 0.25 ml 10 mM $HAuCl_4$ (Sigma Aldrich, 520918) and ice-cold 0.6 ml 10 mM $NaBH_4$ solution (Sigma Aldrich, 71321) was added under vigorous stirring at room temperature for 5 minutes or until the solution color changed to brown from yellow indication reduction of gold salt and formation of gold seed. Next to synthesize the required gold nanorods, 40 ml 0.1 M CTAB, 2 ml 0.01 M $HAuCl_4$ aqueous solution, 0.5 ml 0.01 M $AgNO_3$ (Sigma Aldrich, 20439 0), 0.8 ml 1 M HCl (Sigma Aldrich, H9892), 0.32 ml 0.1 M ascorbic acid (Sigma Aldrich, A92902) and 96 µl of 50 times diluted seed solution were added sequentially. Each addition is followed by gentle homogenization and after last addition the subsequent solution is left in dark for 24 h. Finally, the solution was centrifuged at 7,000 rpm for 30 mins to remove excess reactants and was then reconstituted in nanopure water for further use. Thus, synthesized AuNRs were subsequently employed to synthesize silver cuboids with absorbance wavelength of 650 nm. Briefly, the AuNRs were centrifuged again (two times in total) and redispersed in 0.2 M hexadecyltrimethylammonium chloride (CTAC) (Sigma Aldrich, 52366) solution to complete solvent exchange. The resulting solution was then stored at 60° C for 20 minutes. Thereafter, 150 µl of 10 mM $AgNO_3$ and 150 µl of ascorbic acid was added sequentially and the solution was stored at 60° C for 4 hours.

Conjugation procedures (Cy5-BSA-Biotin):

Employing EDC–NHS chemistry, first BSA, used as the stabilizing agent, was conjugated with biotin, used as universal recognition element. For this, solution of 2.2 ml 5 mg ml−1 BSA (Sigma Aldrich, A7030) in 1X PBS and 2 mg NHS–PEG4–biotin (Thermo Scientific, 21329) was incubated at room temperature for 1 hour. The resulting BSA-biotin conjugates were purified by a desalting column (Thermo Scientific, 21329, 7,000 MWCO). The columns were pre-equilibrated with amine-free 1X PBS solution. Next, to conjugate Cy5 dye to BSA-biotin, 220 µl 1M of $K_2HPO_4$ buffer was added to 2.2 ml of purified BSA-biotin solution. $K_2HPO_4$ is employed to increase the pH of the solution to 9. Subsequently, 25 µl of 5 mg $ml^{-1}$ NHS–Cy5 (Fluorophores, 1501-5) was added to the above solution, and incubated at room temperature for 2 hours. Finally, conjugated BSA–biotin–Cy5 was purified using a Zeba desalting column pre-equilibrated with nanopure water.

Synthesis of on-plasmonic fluors :

Silver cuboids (AuNRAg) was employed as the plasmonic core to prepare plasmonic fluor–Cy5. 1 µl of (3-mercaptopropyl)trimethoxysilane (MPTMS) (Sigma Aldrich, 175617) and 1 ml of AuNRAg (extinction ~2) were mixed and shaken on rocking bed for 1 hour. Next, 2 µl of APTMS (Sigma Aldrich, 281778) and 2 µl of trimethoxy(propyl)silane (TMPS) (Sigma Aldrich, 662275) was added to the MPTMS-modified AuNRAg to form the polymer spacer layer. Excess monomers were removed from the AuNRAg–polymer solution by two centrifugations at 6000 rpm for 10 min. After each wash the pellet was redispersed in 1 M CTAC aqueous solution to ensure colloidal stability. Polymer coated AuNRAgs were concentrated into a final volume of 5 µl. Next, to conjugate Cy5-BSA-Biotin complex to polymer modified AuNRAgs, we followed procedures mentioned in previously reported study[7]. Briefly, to allow coating of Cy5-BSA-Biotin to AuNRAgs, the pH of 100 µl 4 mg $ml^{-1}$ Cy5-BSA-Biotin was lowered by adding 1 µl of 20 mg $ml^{-1}$ citric acid (Alfa Aesar, 36664). To this solution, concentrated AuNRAg-polymer solution was added, and the resulting solution was sonicated for 20 min in dark. After coating, excess Cy5-BSA-Biotin was removed by centrifugation at 3,000 rpm for 10 min and incubated with 0.4 mg $ml^{-1}$ of Cy5-BSA-Biotin in pH 10 nanopure water (1 µl NaOH in 10 ml of water) for 3 days at 4 °C. Finally, the nanostructures were washed 4 times using pH 10 nanopure water by centrifugation at 3,000 rpm for 10 mins. The nanolabels were then redispersed in 1% BSA in 1X PBS solution for use in immunoassays. To further label streptavidin onto these nanolabels, the biotinylated plasmonic-fluor solution was incubated in 100 µg/ml solution of streptavidin for an hour on the rocking bad and subsequently purified by washing 4 times using pH 10 nanopure water by centrifugation at 3,000 rpm for 10 mins.

Synthesis of off resonance plasmonic fluors:

To prepare plasmonic core off resonance to Cy5 dye, first AuNRs with absorbance wavelength 950 nm (localized surface-plasmon resonance wavelength) were synthesized by the seed-mediated method as described above. Thus, synthesized AuNRs were subsequently employed to synthesize silver cuboids with absorbance wavelength of 800 nm. Briefly, the AuNRs were centrifuged again (two times in total) and redispersed in 0.2 M hexadecyltrimethylammonium chloride (CTAC) (Sigma Aldrich, 52366) solution to complete solvent exchange. The resulting solution was then stored at 60° C for 20 minutes. Thereafter, 20 µl of 10 mM AgNO3 and 20 µl of ascorbic acid was added sequentially and the solution was stored at 60° C for 4 hours.

Silver cuboids (AuNRAg) was employed as the plasmonic core to prepare plasmonic fluor–Cy5. Similar procedures were employed to perform coating of polymer spacer layer and for coating of Cy5-BSA-Biotin to synthesize off biotinylated resonance plasmonic fluors. To further label streptavidin onto these nanolabels, the biotinylated plasmonic-fluor solution was incubated in 100 µg/ml solution of streptavidin for an hour on the rocking bad and subsequently purified by washing 4 times using pH 10 nanopure water by centrifugation at 3,000 rpm for 10 mins.

**Supplemental Note 3:**

Collection Efficiency (CE) calculation -

The fluorescent dye molecules were modelled as a dipole source placed in between the plasmonic nanostructure and the PC/glass. The electric field profile was recorded below the grating structure and above the PF structure and projected to the far field with the intensities $I_b(k_x, k_y)$ and $I_a(k_x, k_y)$respectively. $I'_b$ was obtained after applying the Fresnel coefficient to $I_b$ considering the losses encountered by the radiation passing through the glass/air interface. The collection efficiency (CE) was calculated by considering the finite angular bandwidth of the collection objective -

$$\mathrm{CE} = \frac{\iint_{\mathrm{k_x^2+k_y^2<NA}} (\mathrm{I'_b}) \mathrm{dk_x dk_y}}{\iint_{\mathrm{k_x^2+k_y^2<1}} (\mathrm{I_a} + \mathrm{I_b}) \mathrm{dk_x dk_y}}$$

On evaluating the above equation for an NA = 0.25 and the peak fluorescence wavelength of 665 nm, we obtained the CE to be 3.8% for PC substrate compared to 1.92% for the $TiO_2$ substrate, resulting in 2-fold increase in the collection efficiency.

Quantum Yield calculation -

The radiative and non-radiative decay rate can be calculated for Cy5 from the measured lifetime and Quantum yield as [8]

$$k_r = \frac{Q}{\tau} = 0.27\ \mathrm{ns}^{-1}$$

Where $k_r$ is the radiative decay rate, Q (=0.27) is the quantum yield and τ is the measured decay rate (= 1ns). The non-radiative decay rate can be calculated as

$$k_{nr} = \frac{1}{\tau} - k_r = 0.73 ns^{-1}$$

Assuming the $k_{nr}$ is unchanged after the conjugation to the nanostructure (since minimal quenching was observed[9]). The radiative rate enhancement and increase in quantum yield on conjugation with the plasmonic nanostructure can calculated as:

$$k_r^{\mathrm{PF}} = \frac{1}{\tau_{PF}} - k_{nr} = 2.318\mathrm{n}s^{-1}$$

$$Q^{PF} = k_r^{PF} * \tau_{PF} = 0.76$$

Where $\tau_{PF}$ $(=\ 0.328\ ns)$ represents measured lifetime after the conjugation with NP (Fig. 3 (d)). As a result, the quantum yield is improved 2.8-fold (from 27% to 76%) on conjugation with NP.

**Supplementary Note 4: Optimizing surface conjugation of antibodies on the PC surface**.

Two silanes, (3-Aminopropyl)trimethoxysilane (APTES) and (3-Isocyanatopropyl)triethoxysilane and were initially compared to test antibody immobilization efficiency on the photonic crystal surface. The surfaces were washed by sonication in acetone, isopropanol, and MilliQ water, then were oxygen plasma-treated (PicoDiener) and placed into a 5% silane solution in THF for 30 minutes. The photonic crystals were then washed in THF, acetone, ethanol, and MilliQ water, and dried under nitrogen gas. The capture antibodies were added for 1 hour at room temperature at a concentration of 0.2 µg/ml, then the standard DuoSet Human IL-6 ELISA protocol was performed. However, the isocyanate silane was initially too reactive with the detection antibody, causing a high background signal from non-specifically binding PFs. Therefore, the isocyanate silane was used along with an inert blocking silane, in a 95:5 ratio, with butyl(chloro)dimethylsilane, which was found to reduce non-specific binding without effect on standard detection. All silanes were purchased from Sigma Aldrich.

To further increase the efficiency of antibody immobilization, different surface chemistry methods were analyzed through the comparison of a biotinylated capture antibody and a non-biotinylated capture antibody to reveal the streptavidin-coated PF binding efficiency (Fig. S6 A). Additionally, a bifunctional linker molecule, N,N'-Disuccinimidyl carbonate (DSC) was evaluated as a subsequent binding method. The DSC linker acts as a secondary reaction component to link amines in antibody lysine side chains of exposed amino acids. This allows antibody linkage to isocyanate groups that do not undergo rapid conjugation, which are instead converted to amines. The same increase of (2-fold) of immobilized capture antibody is seen in the addition DSC to APTES (Fig. S6)
